# A Systematic Comparison of tTIS Optimization Approaches for Focal Neuromodulation

**DOI:** 10.64898/2026.05.18.726031

**Authors:** Paul Ghanem, Armin Moharrer, Mathew Yarossi, Alan D. Dorval, Dana H. Brooks, Sumientra Rampersad

## Abstract

Transcranial temporal interference stimulation (tTIS) is a promising non-invasive brain stimulation technique that has the potential to selectively modulate deep brain regions by delivering two high-frequency alternating currents that interfere to produce a low-frequency amplitude-modulated envelope at the target. A key challenge in deploying tTIS is finding electrode current patterns that are simultaneously effective, focal, and safe. This is a fundamentally non-convex optimization problem for which a number of methods have recently been proposed. However, no systematic comparison of these methods across a large and diverse set of brain targets has been performed, leaving practitioners without clear guidance on how best to optimize for a particular experimental setting. In this work, we present a comprehensive benchmarking study comparing seven tTIS optimization methods that have appeared in the literature in recent years: exhaustive search, genetic algorithm, multi-objective evolutionary algorithm (MOVEA), min-max optimization, convex TI (CVXTI), non-convex optimization with convex relaxations, and an unsupervised neural network. All methods were evaluated across 250 brain targets spanning cortical and subcortical gray matter and white matter regions in five finite element head models. Each method was evaluated on two key metrics: mean electric field strength within the target region of interest, and off-target stimulated brain volume. Results were stratified by tissue type and target depth to identify systematic performance differences. Based on these results, we provide practical evidence-based recommendations for optimization method selection among these seven methods depending on target location, tissue type, and available computation time. Moreover we provide the code base that will allow other investigators to use these methods for their own applications. Our goal is to provide researchers and clinicians with a clear, evidence-based framework for choosing a tTIS optimization method suited to their specific target and application.

## 1. Introduction

Transcranial current stimulation (tCS) entails sending weak currents through the brain via electrodes on the scalp (generally ≤2 mA per electrode). Both direct (tDCS) and alternating (tACS) currents have been used for over two decades to modulate the activity of superficial cortical areas in basic and clinical research, demonstrating physiological and behavioral effects in areas such as working memory, stroke recovery and visual processing [2, 18, 19, 17].Stimulation devices for tCS are relatively low-cost, small, portable, and induce only minor side effects, but tCS-induced electric fields have poor spatial focality. Relatively high field strengths can occur in large areas surrounding the region of interest (ROI), and also in smaller areas far away[7, 21].While tCS can induce fields strong enough to modulate, although not generally excite, deep neural tissue [14, 4], in such cases superficial tissue closer to the scalp inadvertently experiences larger fields. Transcranial temporal interference stimulation (tTIS) is a relatively novel form of tCS that has the potential to overcome these limitations. Applying tTIS requires two sets of scalp electrodes simultaneously providing alternating currents at nearby frequencies *f*_1_ and *f*_2_ (≥1 kHz) (Fig. 1), which produce an amplitude-modulated signal at carrier frequency *f*_carrier_ = (*f*_1_ +*f*_2_)*/*2 with an envelope that oscillates at beat frequency *f*_beat_ = *f*_2_ − *f*_1_. In mice, tTIS was reported to induce neural firing at *f*_beat_ in deep areas without exciting overlying areas[12].Simulations with finite element models suggested that tTIS-induced electric fields are not strong enough for suprathreshold stimulation in humans, but could achieve subthreshold neuromodulation similar to tACS with greater focality and maximal effects in deep brain areas[22].Initial experiments in healthy and clinical populations have shown tTIS to be safe, well tolerated, and capable of engaging deep brain targets in humans[8].

**Figure 1.**
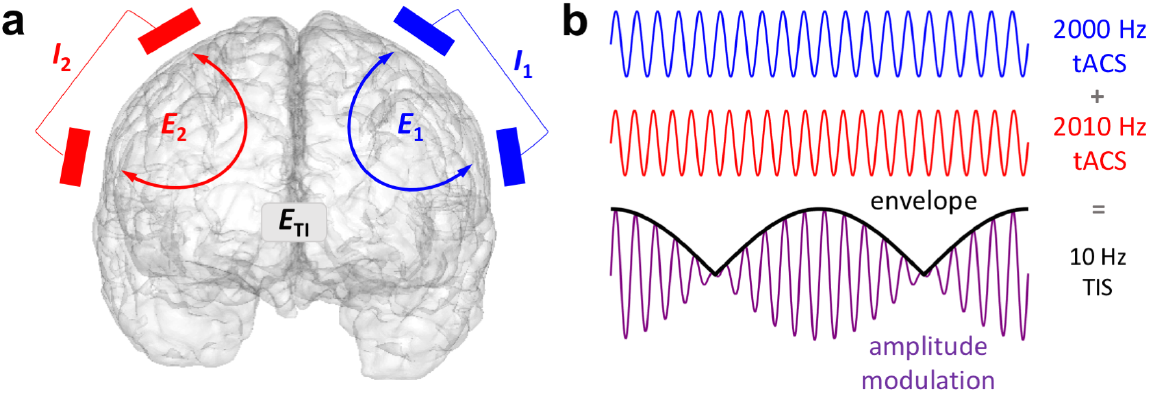
Concept of temporal interference stimulation. **a)** Two pairs of scalp electrodes each supply an oscillating current that produces an electric field in the brain. The intersection of the two fields produces an amplitude-modulated field 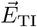 . **b)** Two high-frequency alternating currents add up to one amplitude-modulated oscillation with a carrier frequency equal to the average frequency of the inputs and an envelope oscillating at the difference frequency.

When designing experiments with tTIS, one must select an electrode configuration, or montage, with specified current values. Ideally, this montage is simultaneously *effective* (i.e., reaches sufficient field strength in the ROI for neuromodulation), *focal* (i.e., achieves minimal stimulation of surrounding brain tissue), and *safe* (i.e., stays within accepted current limits). Computational optimization can be used to find a montage that satisfies those constraints. Unlike conventional tCS, the tTIS optimization problem is fundamentally non-convex: the amplitude-modulated envelope arises from the nonlinear interaction of two carrier fields, which prevents straightforward application of linear optimization methods.

There is considerable evidence that neurons respond preferentially to stimulation when the applied electric field is oriented along the predominant linear direction of the neuron or its axon [25, 20].This *preferred direction* can be reconstructed from diffusion tensor imaging (DTI) measurements, but this data is rarely acquired for neuromodulation experiments. In the cortex, the preferred direction can be approximated by calculating vectors perpendicular to the surface, but this does not apply to the deeper brain regions that are often the ROIs for tTIS. We note that the equations describing the tTIS beat field[12]allow us to calculate the field strength in the preferred direction as well as the maximum field strength in any direction, which can be useful when the preferred direction is unknown. However, maximizing over all directions makes the optimization problem yet more difficult to solve.

In recent years, several approaches have been proposed in the literature to address tTIS optimization. The most straightforward, exhaustive search (ES) over all possible combinations of four electrodes in a given set of potential locations achieves good results, but can take days on a PC, or hours on a computing cluster, to optimize for a single ROI[22,16].While using more than four electrodes could increase focality by distributing currents more favorably, or increase efficacy by allowing more current to be safely applied, the computation time using ES optimization would grow exponentially. Genetic algorithms (GA) offer a faster alternative, but existing formulations remain limited to four electrodes and still require substantial computation time[29].A suggested multi-objective evolutionary algorithm (MOVEA) [31] combines GA with multi-objective particle swarm optimization (MOPSO) [6].Several proposed methods extend the electrode count, including a min-max formulation (MinMax)[15],a convex optimization for TI formulation (CVXTI)[11], and a non-convex optimization formulation using convex relaxations (NCOCR)[24], all of which leverage conventional tCS optimization frameworks. Finally, an unsupervised neural network (USNN) has been proposed[1]for TI optimization. In addition to the different formulations and optimization methods, the descriptions of these methods in the literature differ in how many electrodes and how much current can be used and whether they can optimize for the preferred and/or free field direction (see Table 1).In addition, some methods have unique characteristics that may not be practical. For example, the MinMax method allows each electrode to provide stimulation at two frequencies simultaneously, which most actual stimulator systems cannot do. The USNN method optimizes over the white matter surface, which severely restricts the number and location of ROIs it can be used for. Finally, several methods do not have published code, which strongly limits their broader usability.

**Table 1.**
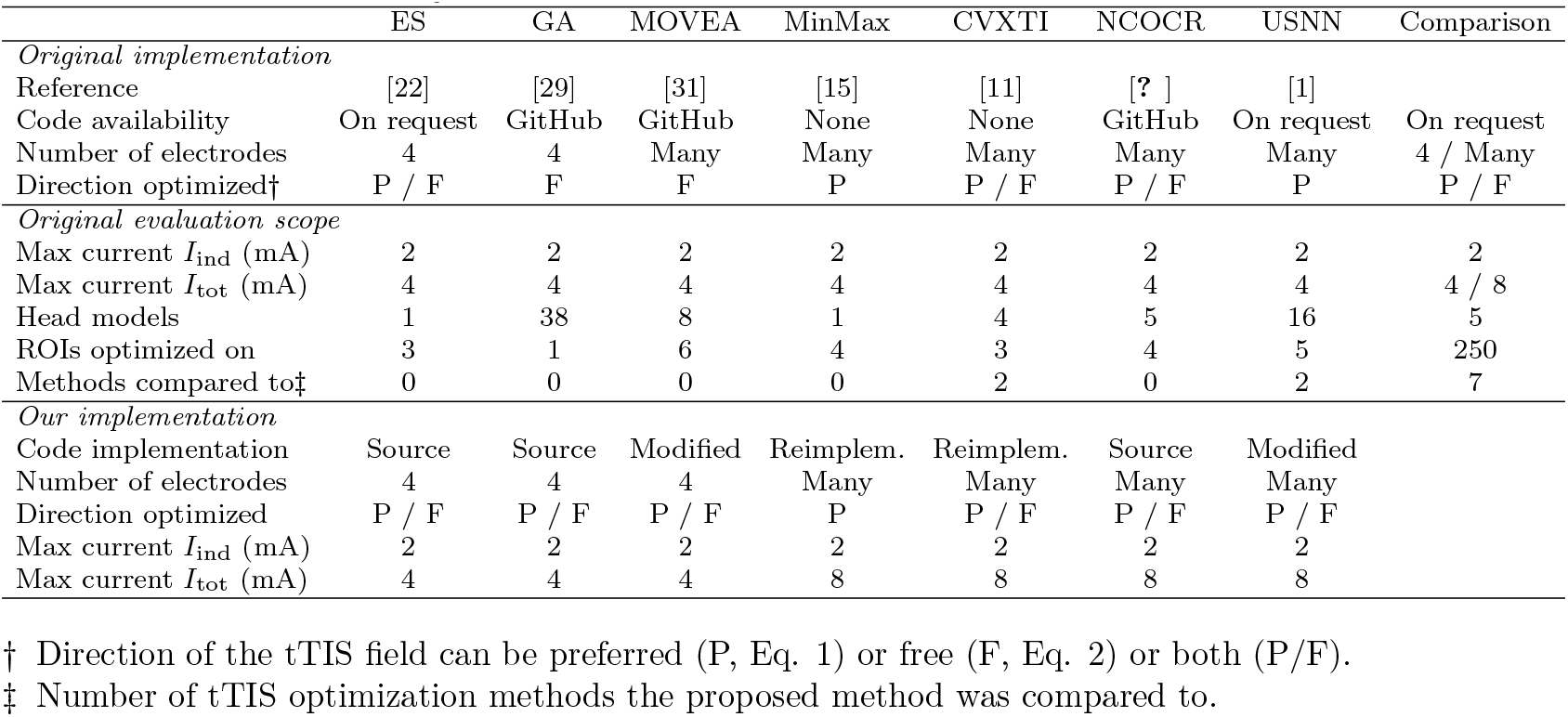
Overview of published tTIS optimization methods and parameters of the current study.

This growing landscape of tTIS optimization methods makes it difficult for practitioners to choose an appropriate approach for a given clinical or research application. When new methods are proposed, they are often evaluated on a small number of ROIs and not compared to the previously published methods. Readers cannot easily compare results across publications themselves, because authors use different head models, locations and shapes of ROIs, and evaluation criteria (see Table 1).For instance, Stoupis and Samaras [29] only tested 1 ROI and compared against a ‘non-optimized’ configuration. Huang et al. [15] used 1 head model and a preferred direction radial to the skull instead of a physiological representation. Bahn et al. [1] compared their USNN method to a two-pair GA, multi-electrode GA, and to Huang et al.’s MinMax method for a single ROI. Geng et al. [11] compared their CVXTI method against MOVEA and MinMax in 3 ROIs. Most authors only compared their results against tACS and not tTIS optimization [22, 29, 31, 15].The studies listed above evaluated their methods using 1–6 ROIs in 1–38 head models and compared to 0–2 competing methods. None examined how performance varied across tissue types or with target depth, both of which are critical factors since tTIS is of interest for ROIs throughout the brain.

In this work, we present a comprehensive comparison study of these seven tTIS optimization methods (ES, GA, MOVEA, MinMax, CVXTI, NCOCR, and USNN) evaluated across 250 brain targets spanning cortical and subcortical gray matter and white matter regions in five different realistic head models. We assess each method on two key metrics: efficacy, specified as mean electric field strength within the ROI, and focality, specified as off-target stimulated volume (percentage of brain volume at or exceeding 0.2 V/m). We evaluate these methods in terms of: 1) the tradeoff between efficacy and focality, 2) how performance scales with target depth and tissue type, and 3) computation time. This comprehensive comparison approach will provide practical guidance on method selection. We also provide all code in an easily usable and freely available format. Our goal is to provide researchers and clinicians with a clear, evidence-based framework for choosing a tTIS optimization method suited to their specific target and application.

## 2. Methods

In this section, we describe head model construction and ROI definitions, simulationa and the tTIS field calculation (Section 2.1),optimization metrics and methods implemented in the current study (Section 2.2),and finally our evaluation framework (Section 2.3).In the descriptions that follow, “the TI field” signifies the electric field at the beat frequency, i.e., the field that is the envelope of the amplitude-modulated field (note that its amplitude is the “modulation depth” of the AM waveform), which will be denoted as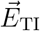.

### 2.1. Modeling and simulation

*Head model construction* Finite element models were created from MRI data of five healthy adults (1 M / 4 F, age 19–27 years) who gave written informed consent prior to scanning. All MRI scans were acquired on a Magnetom Prisma 3T scanner (Siemens, Munich, Germany) with a 64-channel head coil, 0.9×0.9×0.9 mm voxel size, GRAPPA parallel imaging (acceleration factor = 2), and a field of view that captured the complete head. T1-weighted (T1w) images were acquired with an MP-RAGE pulse sequence (TR = 2700 ms, TE = 3.68 ms, TI = 1090 ms, flip angle = 9°) with fat suppression and image dimensions of 224×256×288 voxels. T2-weighted (T2w) images were acquired with an SPC pulse sequence (TR = 3200 ms, TE = 408 ms) and dimensions of 224×256×256 voxels.

The T1w and T2w images were co-registered and segmented using the SimNIBS headreco pipeline (`headreco -all`) [30].Initial segmentation masks were manually corrected in Seg3D [5],resulting in 15k–250k changed voxels across models. The results were reprocessed in SimNIBS headreco to produce triangular surface meshes (`headreco surfacemesh --no-cat`), which were combined into a high-quality 3D Delaunay triangulation via TetGen v1.6.0 [28] (`tetgen -pqA`). This procedure resulted in meshes consisting of 547k–658k nodes and 3.06M–3.74M linear tetrahedral elements with a median element size of 0.49–0.59 mm^3^ and total brain volume of 1172–1439 cm^3^.

All compartments were assigned isotropic conductivity values: skin: 0.465 S/m; bone: 0.021 S/m; eyes: 1.5 S/m; cerebrospinal fluid (CSF): 1.65 S/m; gray matter (GM): 0.276 S/m; white matter (WM): 0.126 S/m. One of the resulting models is shown in Fig. 2A. Although tissue conductivity is frequency dependent and tTIS involves frequencies much higher than those used for tACS, prior tTIS simulations reported qualitatively similar results for standard conductivity values at 0 versus 1 kHz [13].

**Figure 2.**
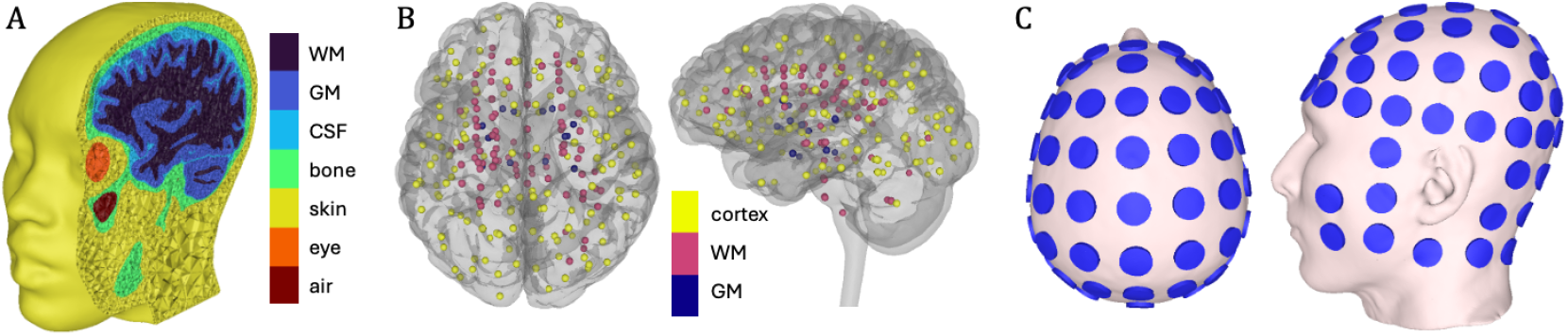
One of the head models used in this study, showing A) the tetrahedral mesh with seven tissue labels, B) the 250 ROI centers colored by ROI type inside a transparent brain surface, C) the 88 electrodes on the skin surface.

For elements in the cortex of the GM compartment, the preferred direction vector 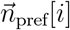 was constructed by interpolating between the nearest normal vectors perpendicular to the GM and WM surfaces in element *i*. Outside of the cortex,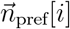 is not defined.

#### Definition of ROIs

Each T1w scan was processed in FreeSurfer 7.4.1 [10] (`recon-all -all -s -i`), resulting in a parcellation mask (`aparc.a2009s+aseg.mgz`) based on the Destrieux brain atlas [9].The mask was split into GM and WM masks before mapping its 176 brain regions onto the respective elements of the head models using SCIRun 5.0 [26] to ensure clean boundaries. For each region, we calculated the volumetric center of the mesh elements in SCIRun. Because there are too few cerebral WM regions in the parcellation for a comprehensive analysis, we constructed a homogeneously spaced grid of 76 ROI centers inside the WM compartment.

For each of 252 ROI centers, we constructed an ROI by growing a sphere from its center until it reached a volume of at least 100 mm^3^ for cortical ROIs or 200 mm^3^ for others, to enforce volumetric similarity across ROIs despite the flat shape of the cortex. ROIs with a radius larger than 10 mm were excluded. The resulting 250 ROIs were grouped into three tissue categories (Fig. 2B): subcortical gray matter (14), white matter (88) and cortex (148), including gyrus (78) and sulcus (76) ROIs. The depth of each ROI was calculated as the Euclidean distance from the ROI center to the nearest point on the scalp surface. Across head models, the ROIs had mean (± std) depths of 39±14 mm, volumes of 145±52 mm^3^, radii of 3.8±0.6 mm, and contained 122±108 elements.

#### Simulations and lead field construction

On the scalp of each head model, we identified 61 locations of the standardized 10-10 system for electrode placement [3, 27]using custom code and manually placed fiducials. We added 27 locations on the neck and cheeks by visual inspection, leading to a total of 88 electrodes covering the entire head minus the face (Fig. 2C). At each location, we constructed a cylindrical electrode with a radius of 10 mm and a height of 3 mm. For 87 electrodes, we solved the Laplace equation ∇·*σ*∇Φ = 0, where *σ* and Φ are the electric conductivity and potential respectively, with a current of 1 mA flowing between that electrode and the 88th electrode, in SCIRun.

We calculated 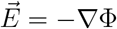 and stacked the resulting electric fields into a lead field matrix *A* of size *N* × 87 × 3, where *N* is the number of brain elements in a model.

#### TI field calculation

Given the current vectors *I*_1_, *I*_2_ ∈ ℝ^*L*^, which contain the current amplitudes of *L* electrodes at frequencies *f*_1_ and *f*_2_, respectively, we can calculate the two carrier fields 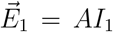 and 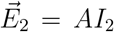 . The TI field at each brain element, *k* ∈ {1, … , *N*}, where *N* is the total number of brain elements, was computed from the carrier fields using different metrics depending on whether a preferred direction was available for that element.

For elements where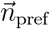 is defined, i.e., the cortex in our models, the TI envelope amplitude along the preferred direction is:

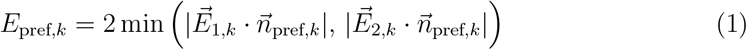

This gives the component of the TI envelope estimated to be most effective at driving neurons oriented along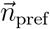 [20, 12].

For elements without a known preferred direction, the TI envelope amplitude is computed as the maximum over all possible orientations. Prior to computation, if the angle *α* between 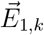 and 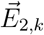 exceeds 90°, 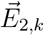 is flipped to ensure *α* ≤ 90°. Denoting the larger- and smaller-magnitude fields as 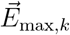 and 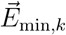 (i.e 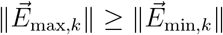) the closed-form solution is [12]:

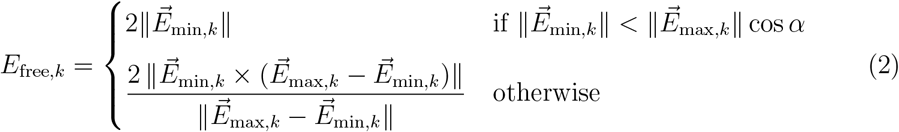

When 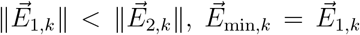 and 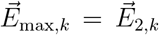 , and vice versa. Note that *E*_free,*k*_ ≥ *E*_pref,*k*_ always, as the free field represents the maximum achievable TI amplitude regardless of direction.

We will optimize the TI field *E*_TI_, i.e., aim to maximize *E*_TI_ in the ROI and minimize it elsewhere, under two conditions. In the *free condition*, we will only consider *E*_free_, so *E*_TI,*k*_ is defined uniformly across all brain elements as:

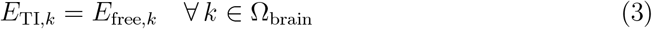

In the *preferred condition*, we aim to maximize *E*_pref_ in the ROI. Since *E*_pref_ is not defined outside the cortex, this will only be done for cortical ROIs. To calculate *E*_TI_ outside the ROI, we will use:

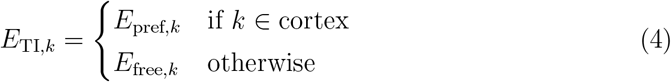

### 2.2. Optimization

Below, we provide a brief description of each optimization method individually; for full algorithmic details we refer the reader to the original publications. Several methods and our evaluation framework use Pareto optimization, which is the process of finding a set of solutions (a Pareto front) that balance two or more conflicting objectives.

All methods were subject to a per-electrode constraint of *I*_ind_ = 2 mA and a total injected current of *I*_tot_ = 4 mA per frequency channel. If any optimized current pattern exceeded either constraint, the currents were scaled down proportionally. For methods that only allow 4 electrodes (ES, GA), the total current is limited to *I*_tot_ = 2 * *I*_ind_. The other methods support multi-electrode configurations that can distribute the higher total current across more electrodes. For these methods, we assessed the effect of increased total current by setting *I*_tot_ = 8 mA (with the same *I*_ind_).

#### Exhaustive Search (ES)

Rampersad et al. [22]evaluate all combinations of two electrode pairs from the set of 88 available electrodes, combined with a range of 21 current ratios between the two pairs. This produces a set of 146M candidate solutions. For each candidate, the median TI field in the ROI and the total stimulated volume are computed, and a Pareto front trading off these two metrics is constructed. We used the original source code provided by the authors and replaced the median TI field with the mean because some of our ROIs have few elements.

#### Genetic Algorithm (GA)

Stoupis and Samaras [29] use a genetic algorithm to search over combinations of two electrode pairs, optimizing both electrode selection and injection currents. The objective function maximizes the electric field ratio between the ROI and surrounding brain tissue, with penalties applied when the ROI field falls below a specified threshold. We used the original source code provided by the authors without modification.

#### Multi-Objective Evolutionary Algorithm (MOVEA)

Wang et al. [31]use a two-stage optimization approach to jointly optimize field strength and focality. First, a genetic algorithm explores the electrode configuration space to identify promising candidate solutions. These are then refined by multi-objective particle swarm optimization (MOPSO), which produces a set of Pareto-optimal solutions. We used the authors’ original implementation with two modifications: the main optimization loop was corrected to prevent an infinite restart condition, and the code was adapted to execute on GPUs to reduce computation time, using NVIDIA V100 accelerators. While MOVEA technically supports multi-electrode optimization, it did not converge when we used *I*_tot_ = 8 mA for any ROI. Hence, we do not report results for this condition for MOVEA.

#### Min-Max Optimization (MinMax)

Huang et al. [15] formulate tTIS optimization as a min-max problem that maximizes *E*_TI,*k*_ over the ROI, i.e., it balances the two current patterns so that the weaker field component is as strong as possible. Off-target field spread is controlled by a quadratic focality constraint that bounds the total modulation depth power outside the ROI below a threshold *P*_max_. All electrodes are shared across both frequencies. The resulting non-convex problem is solved using sequential quadratic programming (`fminimax`), initialized with the corresponding tACS solution. Sweeping *P*_max_ over a logarithmically spaced range traces a Pareto front between target intensity and off-target focality. Since the source code could not be made available, we implemented the method based on the paper. We confirmed correct implementation by comparing our results on the MNI152 head model with those of the author. We then adapted our code to support the 8 mA total current condition.

#### Convex Optimization for TI (CVXTI)

Geng et al. [11] exploit the property of the tTIS envelope that the electric field with the smaller amplitude dominates the optimization: since the TI field is determined by the weaker of the two high-frequency fields, the joint two-pattern optimization can be decomposed into two single-pattern tACS problems, each solvable via convex methods. Electrodes are partitioned into two spatially disjoint groups (e.g., left–right, anterior–posterior, or superior–inferior relative to the target), with one group per frequency. For intensity optimization, each group independently maximizes its target field using a least-squares objective subject to zero-sum and total current constraints; currents of the stronger group are scaled down to match the total current of the weaker group. For focality optimization, the optimization of the first group of electrodes minimizes total brain field power subject to a prescribed target field *E*_0_; the field resulting from the first group of electrodes in a chosen ROI serves as the equality constraint for the second group’s optimization, coupling the two steps sequentially. The electrode pair producing the weaker E-field at the target determines the TI envelope intensity, with the current of the stronger pair scaled down to match. No source code was available for this method; we implemented it from scratch following the algorithmic description in the original publication.

#### Non-Convex Optimization with Convex Relaxations (NCOCR)

Roig-Solvas et al. [24]frame tTIS current pattern selection as a non-convex optimization problem arising from the *E*_TI,*k*_ dependence of the tTIS field on the two input current patterns. To implicitly enforce focality without constraining millions of off-target elements, electrodes are partitioned into two disjoint sets, one per frequency, spatially separating the two high-frequency fields, similarly to CVXTI. For each ROI, *p* such electrode partitions are generated. For each partition, a rank-one approximation of the ROI lead field submatrix renders the objective concave, yielding a convex relaxation that is solved for a global optimum; this solution then initializes a local non-convex solver. The preferred condition thus produces *p* candidate solutions. For the free condition, optimization is repeated along *d* uniformly spaced directions per partition, yielding *d* × *p* candidates. We used the authors’ original code. In our implementation we set *p* = 15 (12 from geometric splitting planes based on normal vectors revolving around the *x, y*, and *z* axes at 0°, 45°, 90°, 135°, plus 3 random assignments with equal probability per electrode) and *d* = 13, yielding 15 × 13 = 195 candidates for the free condition.

#### Unsupervised Neural Network (USNN)

Bahn et al. [1] optimize tTIS electrode currents using a trainable generator network coupled to a fixed, differentiable forward model that encodes the lead field equation and TI envelope computation. The generator maps a unit constant through hidden layers to 2*L* electrode current values (one per electrode per frequency). The forward model contains no trainable parameters; it implements 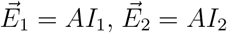 , and the TI field *E*_TI,*k*_ as fixed-weight layers, serving solely to make the physics differentiable for backpropagation. Training minimizes a composite loss combining the peak ratio (target vs. non-target peak modulation), concentration ratio (target vs. whole-brain modulation density), and mis-stimulation ratio (non-target area exceeding the target mean). No labeled data are required. The authors’ implementation used a lead field defined only on the WM surface and consequently required ROIs to be on this surface. We initially projected our ROIs onto the WM surface, but as can be expected, this did not produce usable results for ROIs distant from that surface. Hence, we adapted the method to operate on volumetric lead fields by using a lead field defined over the full finite element mesh to enable optimization for ROIs at any location in the brain.

### 2.3. Evaluation protocol

#### Performance metrics

To compare the performance of all optimization methods, we need to evaluate their solutions using the same metrics of efficacy and focality. A threshold of *E*_th_ = 0.2 V/m was used to define effective neuromodulation. This value is based on estimates of the minimum electric field strength required to modulate neuronal activity in computational and in vitro studies [21, 23]. Efficacy will be represented by *E*_ROI_, the mean TI field strength within the ROI:

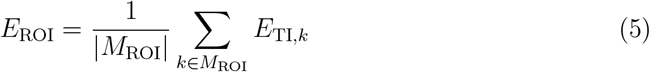

where *M*_ROI_ is the set of elements in the ROI. A method is considered to have successfully stimulated a given ROI if *E*_ROI_ ≥ *E*_th_. Focality will be quantified as *V*_off_, the percentage of total brain volume outside the ROI where the TI field exceeds *E*_th_:

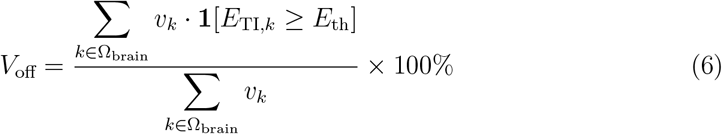

where *v*_*k*_ is the volume of element *k* and Ω_brain_ is the full brain domain.

#### Focality versus intensity conditions

Two evaluation conditions were considered based on different requirements an experimenter may have when choosing an electrode montage for their study. Under the *maximum intensity* condition, we determine the highest ROI field strength a method can achieve while remaining within safety limits. For this condition, only the safety constraints were enforced. Under the *maximum focality* condition, we determine the lowest amount of unwanted stimulation a method can achieve in order to reach effective field strengths in the ROI. For this condition, electrode currents in each solution were uniformly scaled so that *E*_ROI_ = *E*_th_, enabling direct comparison of *V*_off_ across methods at a standardized field strength.

Several of the methods (ES, MOVEA, MinMax, NCOCR) produce multiple solutions per ROI, so we needed to choose which solutions to use for evaluation. For each candidate, *E*_ROI_ and *V*_off_ were computed. A Pareto front was constructed by binning candidates by their *E*_ROI_ value in steps of 0.05 V/m. For the maximum focality condition, we selected the solution with minimal *V*_off_ among all bins where *E*_ROI_ ≥ *E*_th_. For the maximum intensity condition, we selected the bin with the highest *E*_ROI_.

#### Field direction

Not all methods natively support optimization of both field directions (see Table 1).The published source code for GA and MOVEA only supports optimizing *E*_free_ and we implemented *E*_pref_. The USNN source code only supports *E*_pref_ and we implemented *E*_free_. MinMax was designed for *E*_pref_ only and could not easily be extended to support *E*_free_. Hence, we optimized MinMax only for *E*_pref_ and used the solution (i.e., current values) from that optimization to evaluate both the free and preferred conditions. All other methods were optimized separately under both conditions.

#### Statistical analysis

All metrics were computed separately for each subject and averaged across the five subjects per ROI. For a given ROI, if *E*_ROI_ *< E*_th_ for any of the five subjects, the averaged results for this ROI were excluded from comparisons. Results were stratified by tissue type (cortex, white matter, subcortical gray matter) and evaluated with respect to depth of the ROI.

## 3. Results

Across 5 head models, 7 methods were evaluated spanning 250 cortical, white matter, and subcortical gray matter targets under four conditions: maximum focality and maximum-intensity conditions, each with preferred direction and free direction constraints.

### 3.1. Off-target stimulated volume

Figure 3 shows the distribution of *E*_ROI_ under the two maximum intensity conditions and *V*_off_ under the two maximum focality conditions, pooled over all head models and ROIs.

**Figure 3.**
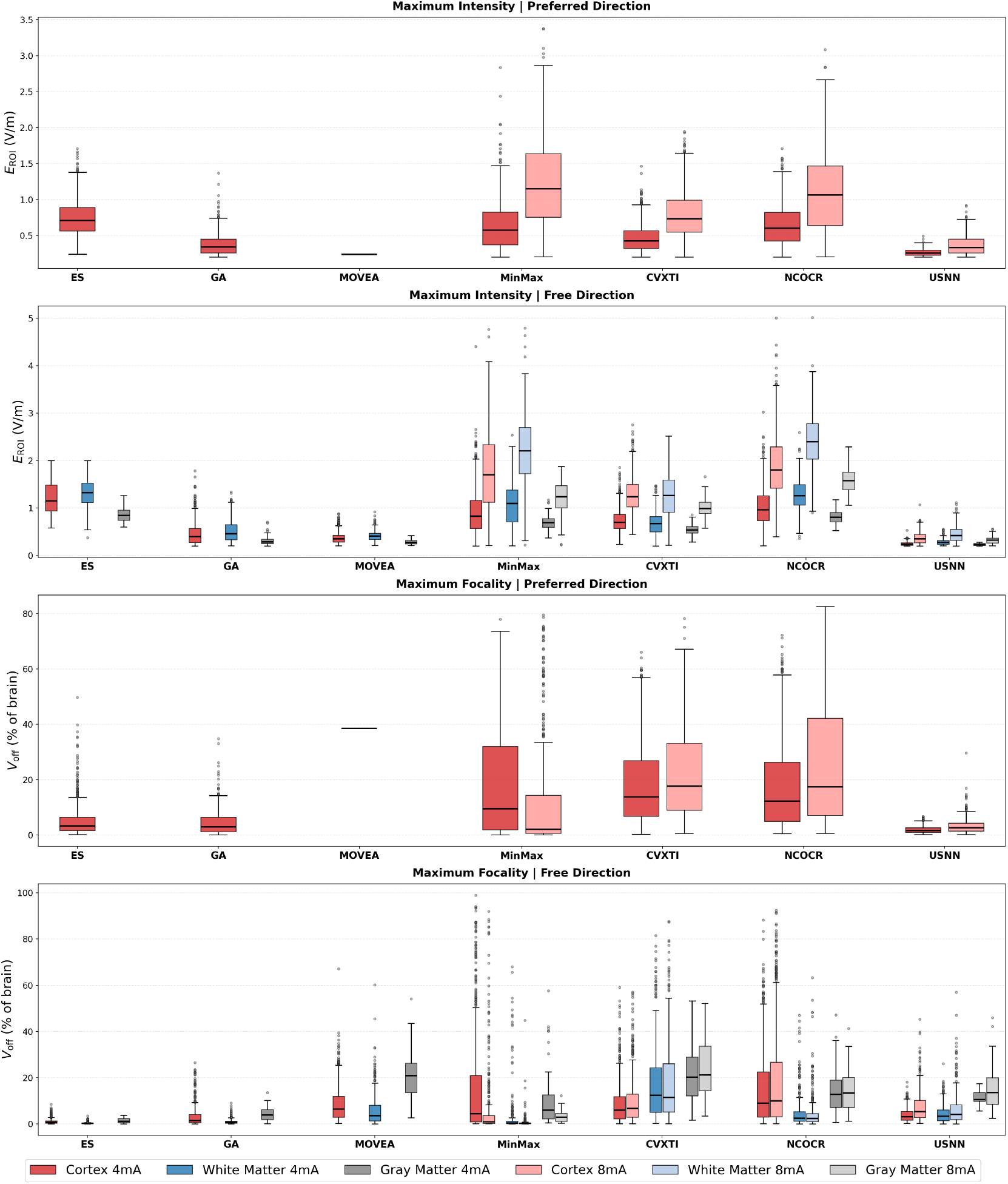
Boxplot distribution of E_ROI_ and *V*_off_ for seven methods across 250 ROIs averaged over five head models. E_ROI_ (mean TI field in ROI) represents efficacy (higher is better) and *V*_off_ represents focality (off-target stimulated volume; lower is better). Boxes span the 25th to 75th percentiles of ROI values with the median marked as a horizontal line. Whiskers extend to the most extreme data point within 1.5 times the box height from the box edges; points beyond are shown individually.

#### Maximum-intensity condition

In this condition, we compare target intensity for each method’s maximum field achievable within the safety constraints. For the preferred field direction (only cortical targets) at 4 mA, ES, MinMax and NCOCR achieved median *E*_ROI_ values above 0.5 V/m. Median values for CVXTI and GA were between 0.25 and 0.5 V/m, with USNN at approximately 0.2 V/m. MOVEA showed near-zero median *E*_ROI_. In the preferred direction condition, MOVEA’s boxplot collapses to a single line because only one ROI passes the threshold. The highest value for individual ROIs was 2.8 V/m achieved by MinMax. In the free direction condition, cortical median *E*_ROI_ exceeded the preferred direction median for all methods except USNN, for which the two were approximately equal. The median for ES exceeded 1.0 V/m for cortical and WM targets. MinMax and NCOCR exceeded 1.0 V/m for WM targets, while their cortical median *E*_ROI_ was approximately 0.8–0.9 V/m. CVXTI occupied intermediate median *E*_ROI_ values across all tissue types. GA and MOVEA showed lower values, with similar median *E*_ROI_ across all tissue types (between 0.4 and 0.6 V/m). USNN reached the lowest values overall; its highest value across all ROIs was around 0.3 V/m. Regarding tissue types, WM median *E*_ROI_ exceeded cortical and GM for all methods except CVXTI. GM median *E*_ROI_ is below cortex and WM for all methods.

Doubling the available current (only possible for MinMax, CVXTI, NCOCR and USNN) caused a 1.5–2-fold median *E*_ROI_ increase across all methods, target types, and direction conditions. For MinMax and NCOCR, median *E*_ROI_ exceeded 2.0 V/m for WM targets, and some cortical and WM ROIs exceeded 4 V/m. The tissue ordering of median *E*_ROI_ remained the same between 4 mA and 8 mA.

#### Maximum-focality condition

In this condition, we compare focality for each method’s most focal yet effective solution (i.e., *E*_ROI_ = 0.2 V/m). ES, GA, MinMax and USNN produced the lowest *V*_off_ (i.e., were more focal) with median values below 10% for cortical targets in both direction conditions, but MinMax showed much larger variability. Its maximum is over 70% for preferred and near 100% for free direction, while the other three methods remain under 50% in all targets and conditions. In the preferred direction condition, MOVEA’s boxplot collapses to a single line because only one ROI passes the threshold. CVXTI and NCOCR produced similar results in terms of both median values and (comparatively high) variability. In the free condition, median volumes for cortical targets were lower than those for preferred for all methods except USNN. The median *V*_off_ ranged from approximately *<* 1% to *>* 22% across methods and ROI types, with outlier *V*_off_ values of *>* 60% for MOVEA, MinMax, CVXTI, and NCOCR. For CVXTI and USNN, WM targets produced higher medians than cortical targets; for the other methods, results were slightly lower. All methods produced the largest medians for GM targets. For cortical targets (preferred and free) and GM targets, doubling the available current to 8 mA (only possible for MinMax, CVXTI, NCOCR and USNN) slightly increased median *V*_off_ for all methods except MinMax. MinMax improved focality at 8 mA compared to 4 mA across conditions and ROI types, suggesting that its optimization objective benefits strongly from the additional current budget. Doubling the current produced minor changes for WM targets.

### 3.2. Depth dependence of stimulated volume and mean TI field

Figure 4 shows *E*_ROI_ and *V*_off_ for individual ROIs as a function of ROI depth. Each point represents the average across five head models. Cortical and subcortical gray matter ROIs were combined into one subplot because results were similar.

**Figure 4.**
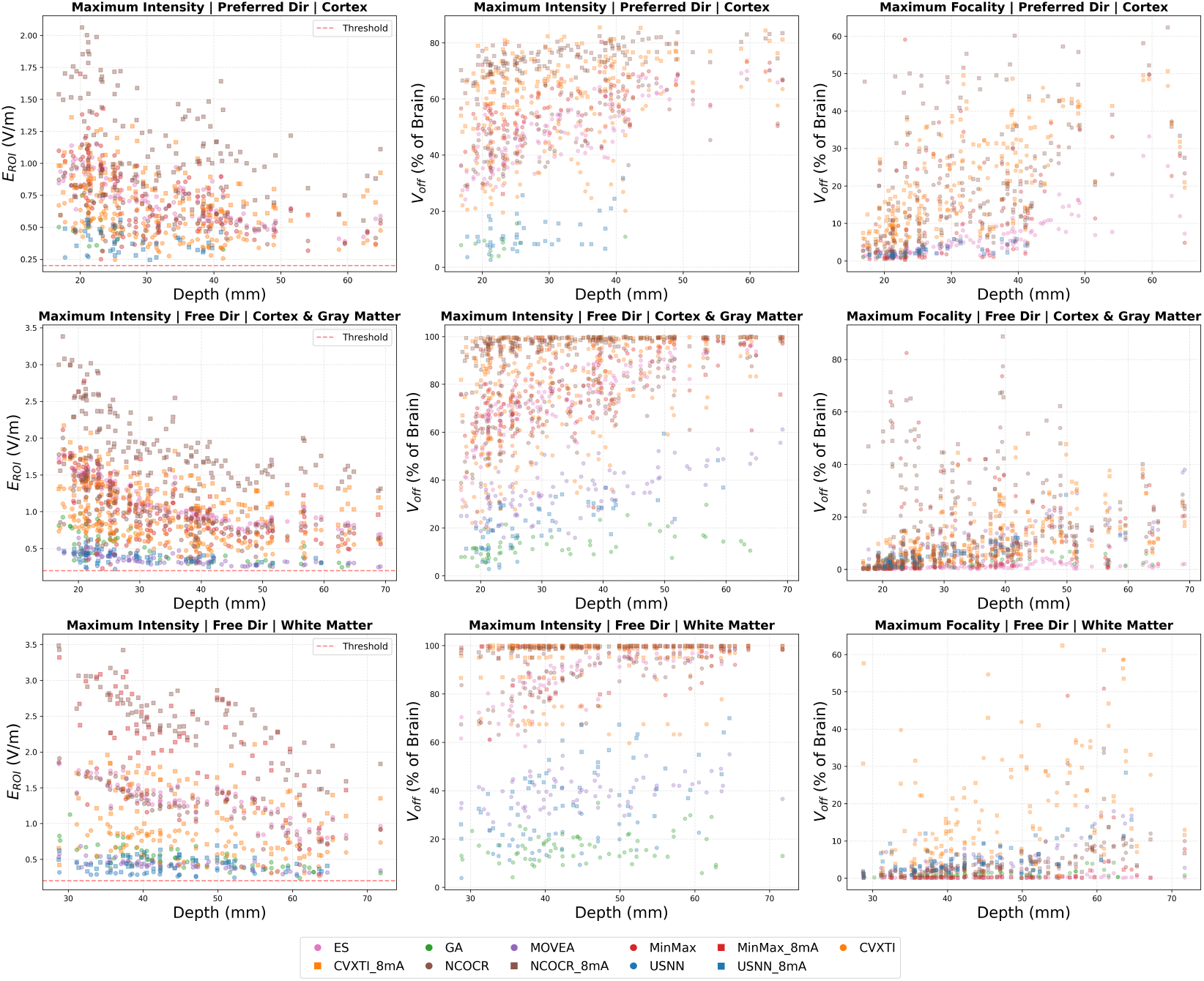
Scatter plots of *E*_ROI_ and *V*_off_ as a function of ROI depth for all targets. *E*_ROI_ (mean TI field in ROI) represents efficacy (higher is better) and *V*_off_ represents focality (off-target stimulated volume; lower is better). Each point represents results for one ROI averaged across five subjects. ROIs that did not pass the threshold for successful neuromodulation (*E*_ROI_ = 0.2 V/m, marked with a horizontal red line) for any of the five subjects were excluded. Circles: 4 mA total current; squares: 8 mA.

#### Maximum-intensity condition

The *E*_ROI_ values generally decreased with depth for all methods, conditions and ROI types (Fig. 4, first column). In cortical and GM targets, NCOCR at 8 mA produced the highest *E*_ROI_ at all depths (peaking above 2.0 V/m in the preferred, and 3.0 V/m in the free condition, for shallow targets), followed by CVXTI at 8 mA and NCOCR at 4 mA. CVXTI and USNN produced the lowest *E*_ROI_ values for the preferred direction, and MOVEA and USNN for the free direction. In WM targets, MinMax and NCOCR at 8 mA produced the highest *E*_ROI_ across all depths, and USNN the lowest.

The *V*_off_ values generally increased with depth for all methods (Fig. 4,second column). In both preferred and free conditions, and for all ROI types, GA and USNN produced the lowest *V*_off_ across the full depth range. These methods also produced low *E*_ROI_. CVXTI and NCOCR at 4 and 8 mA showed the highest volumes in both free and preferred cases for all ROI types, which also matches their high *E*_ROI_ values. MOVEA failed to reach *E*_th_ except in one ROI in the preferred case , and MinMax at 8 mA did not meet the threshold in most cortical ROIs. In targets for which MinMax met *E*_*th*_, stimulation volumes were moderately high compared to other methods for cortical and GM targets, and its 8 mA variant produced the highest volumes in WM targets. Both preferred and free conditions produced large differences between competing methods, almost clustering each method in a distinct *V*_off_ range.

#### Maximum-focality condition

In this condition, *E*_ROI_ is equal for all ROIs and methods by definition, so we only investigate *V*_off_. As in the maximum-intensity conditions, stimulated volumes generally increased with depth for all methods. In the preferred direction, ES maintained the lowest *V*_off_ across most ROI depths, with USNN competitive for shallower targets up to approximately 30 mm. In the free condition, ES produced the lowest *V*_off_ across the full depth range for cortical and GM targets, while MinMax at 8 mA produced the lowest volume in WM targets for most ROIs. In both the preferred and free conditions, CVXTI and NCOCR at 4 and 8 mA showed the highest volumes. In WM targets, NCOCR’s volumes dropped compared to cortical and GM targets, and CVXTI produced the highest volumes. MinMax at 8 mA and MOVEA exceeded *E*_th_ in cortical and GM targets in the free condition, but they did not reach *E*_th_ in most ROIs in the preferred case. Furthermore, MinMax at 8 mA did not reach *E*_th_ in most cortical and GM targets; it produced the lowest volumes across WM targets when the threshold was met. Differences between methods were smaller than for the maximum-intensity condition, because the maximum *E*_ROI_ and overall fields in that condition are vastly different across methods.

### 3.3. Winner analysis

In the previous analyses, we investigated patterns in performance of different methods across ROIs. Here, we will define a metric to choose the “best-performing method” according to that metric for each individual ROI and determine how often each method comes out on top.

#### Maximum-intensity condition

If an experimenter were to desire the highest possible field strength in their target, they would choose a method based on the maximum-intensity condition. However, they may also desire a high (yet not necessarily maximal) field strength that produces a relatively focal field. In order to balance these two requirements, methods were scored using a composite z-score metric:

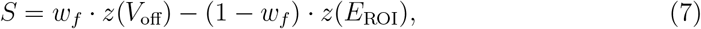

where *z*(·) denotes z-score normalization across methods for that ROI, *w*_*f*_ ∈ [0, 1] is the focality weight, and 1 − *w*_*f*_ is the complementary field strength weight. A lower score indicates a better method: positive *z*(*V*_off_) penalizes high stimulated volume (low focality), while negative *z*(*E*_ROI_) rewards high target field strength. The method with the lowest *S* is declared the winner for each ROI. Figure 5 shows the percentage of ROIs won by each method as a function of *w*_*f*_ , applied to maximum-intensity fields with preferred and free direction ROIs pooled.

**Figure 5.**
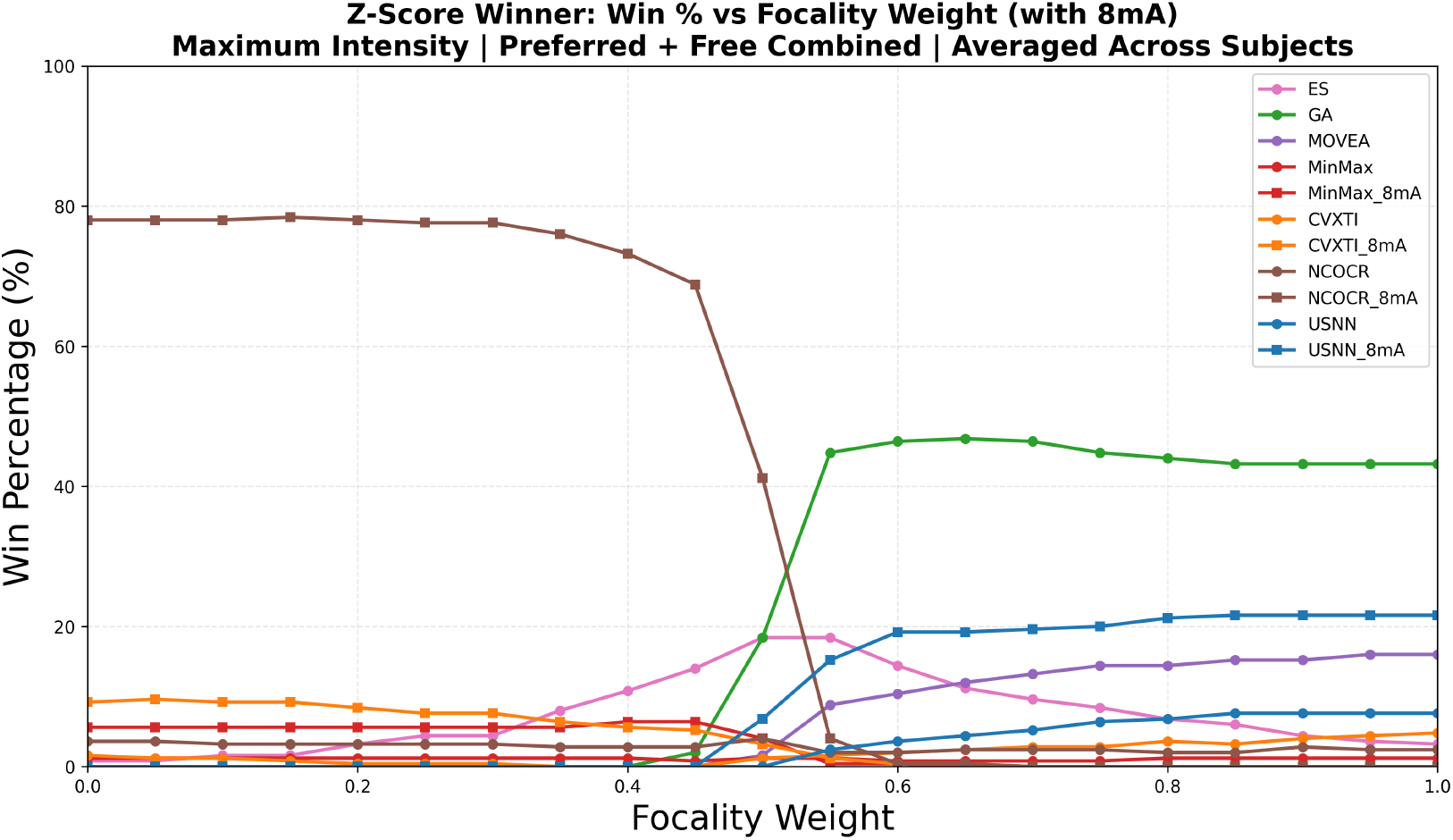
Percentage of ROIs for which each method performed best as a function of the focality weight *w*_*f*_ under the maximum-intensity condition. Performance was quantified using a composite z-score metric that weights efficacy and focality. Results for preferred and free direction conditions were pooled. Circles: 4 mA total current; squares: 8 mA.

When field strength was the sole or primary objective (*w*_*f*_ ≤ 0.45), NCOCR 8mA was the dominant winner, consistently capturing between 70% and 80% of ROIs across the range of weights. This reflects the higher injected current and resulting field amplitudes of this method across all target depths. A sharp transition occurred as focality weight increased past 0.45. NCOCR 8mA win percentages dropped precipitously to below 1% and remained there for *w*_*f*_ ≥ 0.6, while GA emerged as the dominant winner for *w*_*f*_ ≥ 0.55, winning approximately 45% of ROIs. USNN 8mA, MOVEA and USNN 4mA rose concurrently, winning approximately 20%, 15% and 10%, respectively. This transition indicates that GA, USNN and MOVEA achieve a favorable combination of moderate field strength with substantially lower stimulated volume compared to MinMax, CVXTI and NCOCR at both 4 and 8 mA. While all methods show either a sharp transition between two zones or a flat profile, ES peaks for intermediate weights, with winning percentages rising to almost 20% up to *w*_*f*_ = 0.5, and falling back gradually from *w*_*f*_ = 0.55. Notably, CVXTI 8 mA), NCOCR at (4 and 8 mA) and MinMax won zero ROIs when *w*_*f*_ *>* 0.6.

#### Maximum-focality condition

In this condition, all *E*_ROI_ are the same, so there is no trade-off between stimulation intensity and focality. Figure 6shows the percentage of ROIs for which each method achieved the minimum *V*_off_ (combining preferred and free direction conditions). ES was the most frequent winner, achieving the lowest *V*_off_ in 61.2% of ROIs. GA was second at 13.2%, followed by Minmax at 8mA (10.4%) and 4 mA (6.4%) and USNN 8mA (5.6%). Other methods won less than 5% of ROIs, indicating that these methods were rarely the most focal option.

**Figure 6.**
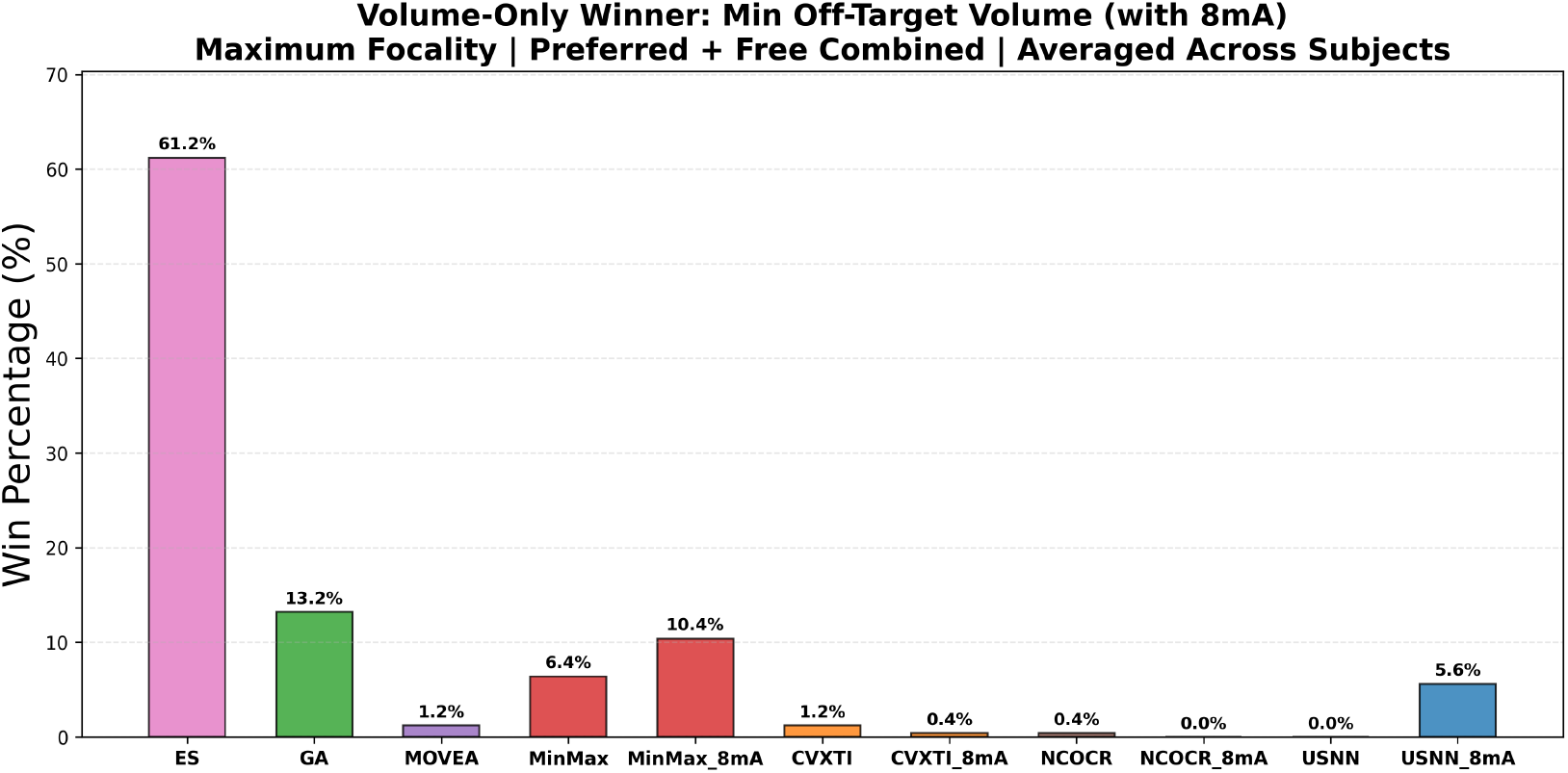
Percentage of ROIs for which each method achieved the minimum stimulated volume under the maximum-focality condition (*E*_ROI_ = 0.2 V/m), pooling preferred and free direction ROIs.

### 3.4. Computational efficiency

Table 2 reports execution times per ROI across 250 ROIs for each optimization method, measured on a single head model. All methods were benchmarked on the same computing infrastructure, and times reflect the complete optimization pipeline including post-processing.

**Table 2.** Execution time per ROI for each optimization method for a single head model. All values are in seconds.

| Method | mA | Mean | Std | Min | Max |
| --- | --- | --- | --- | --- | --- |
| CVXTI | 4 | 69 | 8 | 59 | 84 |
| CVXTI | 8 | 69 | 3 | 59 | 76 |
| MOVEA | 4 | 102 | 55 | 5 | 200 |
| ES | 4 | 467 | 217 | 209 | 1,743 |
| NCOCR | 8 | 824 | 186 | 549 | 1,473 |
| NCOCR | 4 | 950 | 237 | 632 | 2,174 |
| USNN | 8 | 1,636 | 423 | 681 | 3,197 |
| USNN | 4 | 1,741 | 586 | 708 | 3,633 |
| GA | 4 | 25,658 | 20,385 | 874 | 81,379 |
| MinMax | 8 | 62,196 | 30,762 | 7,224 | 147,618 |
| MinMax | 4 | 72,252 | 35,700 | 9,210 | 153,426 |

Computational cost varied by nearly four orders of magnitude across methods. CVXTI was the fastest, completing an ROI in 69 s on average at both current levels with minimal variance (*σ* ≤ 8 s). This predictability is characteristic of convex solvers, where runtime depends on problem dimensions rather than solution landscape complexity. MOVEA was comparable at approximately 102 s per ROI, though with greater variability across ROIs.

ES averaged 467 s per ROI. Its runtime is governed by the number of electrode combinations evaluated and will scale with the electrode montage size (88 in this study). We used 70 cluster nodes simultaneously to achieve these results. Running ES on one CPU would change the mean time from approximately 8 minutes to 9 hours.

NCOCR averaged 15.8 min at 4 mA and 13.7 min at 8 mA per ROI. This includes both optimization across electrode partitions and the subsequent Pareto selection step. The moderate variance (*σ* ≈ 3.5 min) reflects differences in convergence behavior across partitions and ROI locations, though the runtime remains substantially more predictable than the stochastic methods.

USNN required ∼29 min at 4 mA and ∼27 min at 8 mA per ROI, reflecting the overhead of training a dedicated neural network for each target. The similar runtimes across current levels indicate that the network architecture and training procedure, rather than the constraint bounds, dominate computational cost.

GA averaged 7.1 hr per ROI with substantial variance (*σ* ≈ 5.7 hr), ranging from under 15 minutes to over 22 hours. This variability reflects the stochastic nature of genetic algorithms, where some ROIs require substantially more generations to converge than others.

MinMax was the most expensive method, requiring on average 20.1 hr at 4 mA and 17.3 hr at 8 mA per ROI, with a maximum of 42.6 h. Each ROI undergoes multiple iterations of the `fminimax` solver across a range of Pareto sampling points, and the solver is prone to slow convergence near tight constraint boundaries. These runtimes present a practical barrier for multi-subject deployment without aggressive parallelization.

In summary, CVXTI and MOVEA offer per-ROI runtimes on the order of 1–2 minutes, making them practical for iterative workflows and large-scale studies. ES is also fast at 8 minutes per ROI, but this can only be achieved on a computing cluster. NCOCR and USNN require approximately 15 and 28 minutes per ROI, respectively. GA and MinMax require hours per ROI, with MinMax averaging over 17 hours, presenting a potential computational bottleneck for studies with many subjects or ROIs. The variability is also high, with maximum runtimes of nearly 23 hours (GA) and 43 hours (MinMax).

## 4. Discussion

### 4.1. Method ranking and practical recommendations

The results of this comparison reveal that no single tTIS optimization method dominates across all evaluation conditions. The choice of method depends critically on whether the practitioner prioritizes field strength, focality, or a balance of both, and whether a preferred field direction is known. Additionally, results vary strongly across ROIs.

Under the maximum-focality condition, where all methods are compared at a standardized target field strength of 0.2 V/m, ES achieved the highest focality (61.2%) followed by GA(13.6%), MinMax at 8 mA and 4 mA (10.4% and 6.4%) ,and USNN at 8 mA (5.6%). The z-score tradeoff analysis under the maximum-intensity condition provides complementary insight into how method rankings shift when both focality and field strength are considered simultaneously. If maximizing field strength is the sole objective, NCOCR at 8 mA wins approximately 80% of ROIs reflecting the direct advantage of higher injected current. As focality is progressively weighted, GA becomes the dominant method, winning over 45% of ROIs for *w*_*f*_ ≥ 0.55. At the highest focality weights, USNN (at 4 and 8 mA) emerges as the second strongest method, winning up to 30% of ROIs.

Based on our findings, we offer the following practical recommendations. For applications where focality is the primary concern and computational cost is not a constraint, ES, GA,MinMax and USNN at 8 mA are strong choices. When computation time matters, ES provides the best overall balance of focality and field strength across the widest range of trade-off settings, despite being limited to four electrodes. USNN is a strong alternative when maximum focality is desired, particularly given its fast computation time (minutes on a GPU) and competitive performance at high focality weights. MinMax at 8 mA offers a strong focality advantage under maximum-focality conditions. NCOCR and CVXTI at standard current levels (4 mA) are not recommended when focality is a priority, as they were consistently outperformed by other methods in both the normalized and maximum-intensity conditions.

### 4.2. Focality versus intensity tradeoff

The z-score winner analysis (Figure 5) reveals a striking feature of the tTIS optimization landscape: the transition between field-strength-dominated and focality-dominated regimes is abrupt rather than gradual. Below *w*_*f*_ ≈ 0.4, NCOCR at 8 mA and CVXTI together capture ∼95% of ROIs; similarly, above *w*_*f*_ ≈ 0.6, GA, MOVEA and the two USNNs together capture ∼95%. This sharp crossover suggests that the methods occupy distinct niches in the focality-intensity space, with relatively little overlap in the intermediate regime.

This structure has practical implications. For most clinical and research applications, the desired operating point lies in the balanced or focality-leaning range (*w*_*f*_ ≥ 0.5), where the goal is to deliver sufficient field strength to induce neuromodulation while limiting off-target effects. In this regime, GA and USNN dominate, and the choice between them depends on whether the practitioner values moderate field strength with good focality (GA) or maximum focality with lower but still adequate field strength (USNN).

The finding that CVXTI and NCOCR at 4 mA win essentially zero ROIs across all weight settings in the combined maximum-intensity condition warrants discussion. The Two methods do incorporate focality objectives, but their off-target volumes remain higher than those of GA and USNN across this diverse set of targets.

A key observation is that the best-performing methods for focality — ES, GA, MinMax, and USNN — all either directly evaluate, optimize against, or have been trained on the true TI envelope field, whereas the consistently underperforming methods (NCOCR and CVXTI) rely on convex relaxations or approximations of it. This distinction appears more explanatory than any categorization by optimization paradigm, since the top focality performers span sampling-based (ES, GA), continuous minimax (MinMax), and learned (USNN) approaches. Convex decompositions and relaxations used by NCOCR and CVXTI may fail to capture the nonlinear structure of the TI envelope, particularly for deep targets surrounded by tissue with varying conductivity, leading to electrode configurations that are suboptimal in terms of off-target suppression.

MOVEA represents a partial exception: it does evaluate the true TI envelope and performs reasonably well in the free direction condition, for which it was originally designed. However, the preferred direction evaluation is an adaptation of the original implementation, and the evolutionary optimization struggles under this modified constraint, with 97% of ROIs failing to reach *E*_th_. This directional sensitivity limits its practical utility for cortical targets where the preferred direction constraint is typically applied.

### 4.3. Effect of increased current (4 mA vs 8 mA)

Doubling the total injected current from 4 to 8 mA produced method-dependent effects on both field strength and focality. NCOCR at 8 mA achieved the highest maximum field strengths of any method at all target depths, and CVXTI at 8 mA similarly improved over its 4 mA counterpart. However, this increased target field strength represented an overall increase in field strength across the brain, and the focality cost was substantial.

In contrast, MinMax at 8 mA produced lower off-target volumes than MinMax at 4 mA, becoming the top winner in the volume-only comparison. USNN at 8 mA similarly remained among the more focal methods. These results indicate that the effect of increased current is highly method-dependent: for NCOCR and CVXTI it improves field strength at the cost of focality, whereas for MinMax and USNN it yields a net focality advantage or at worst a neutral effect.

In the z-score tradeoff analysis, NCOCR at 8 mA dominated only in the regime where field strength was overwhelmingly prioritized (*w*_*f*_ *<* 0.4). CVXTI at 8 mA did not win at any weight setting. For applications requiring focal stimulation, increasing current for NCOCR or CVXTI does not confer a meaningful advantage, and the increased off-target stimulation may be undesirable from a safety perspective.

### 4.4. Limitations and future work

Several limitations of the present study should be acknowledged. First, all evaluations used a single neuromodulation threshold of 0.2 V/m for the focality metric. This value is commonly used in the tCS literature but represents an estimate of the minimum field strength required for subthreshold neuromodulation. The relative ranking of methods may shift at different thresholds, and future work should examine sensitivity to this parameter.

Results were averaged across five subjects per ROI label. While this captures the central tendency, it does not fully characterize inter-subject variability. A subject-specific analysis would be valuable for clinical applications, especially for certain clinical populations that may have changes in head or brain geometry. We also did not evaluate performance in the preferred direction in the white matter compartment because this data was not available in our models.

Finally, not all methods had source code available. We implemented these methods based on the equations and descriptions in the papers, but there was not always sufficient detail to ascertain that our implementation is identical to those of the authors. We verified results to the best of our ability by repeating analyses in those papers using the same methods, head models and ROIs where possible, before continuing with our evaluation. This work demonstrates the importance of code being published alongside papers.

In terms of methodological extensions, future work could explore adaptive electrode partitioning strategies for NCOCR that are informed by ROI location, hybrid approaches that use fast methods (e.g., USNN or GA) to initialize slower but potentially more accurate optimizers (e.g., NCOCR or MinMax), and systematic comparisons at higher electrode counts to determine whether the focality advantage of multi-electrode methods can be realized without sacrificing computational tractability.

## 5. Conclusion

No single published tTIS optimization method consistently performs better than others and there is large variability across ROIs in any evaluation condition. Recommendations of which method to use depend on an experimenter’s requirements of intensity, focality, direction, and the location and type of their brain target. This work demonstrates the importance of authors of computational papers evaluating their methods across comprehensive sets of models and brain targets, comparing against existing methods, and making their code available online.

